# EVRCEPT: EV RNA Cargo Enrichment Prediction Tool to predict enrichment of RNA into Extracellular Vesicles

**DOI:** 10.64898/2025.12.08.693008

**Authors:** Ahmed Abdelgawad, Jaysheel D. Bhavsar, Shawn W. Polson, Vijay Parashar, Mona Batish

## Abstract

Extracellular vesicles (EVs) are small membrane-bound vesicles that are released by most cells. EVs have been shown to transport molecules including proteins and various types of RNAs between cells of even different types. Furthermore, EV RNAs are shown to modulate gene expression in physiological and pathological conditions in recipient cells which can be utilized in therapeutics by engineering cells to enrich RNA of interest in EVs. However, how specific RNA species are enriched in EVs is a long-standing question in the field. Here, we used sequence features of RNAs to predict its enrichment in EVs. These features include length, nucleotide and dinucleotide frequencies, secondary structure information, number of exons, coding probability for non-coding RNAs as well as RNA binding protein (RBP) motifs. The model achieved a performance (AU-ROC: 90%, 77%) for circRNAs and mRNAs, respectively. Here, we present a web tool called, EV RNA Cargo Enrichment Prediction Tool (EVRCEPT), that allows users to predict likelihood of input RNA to be enriched into EVs. This tool will also provide the list of RBPs that are likely to interact with the input RNA and works with both linear and circular RNAs. This webtool, which is freely accessible at https://euler.dbi.udel.edu/evrcept, will help understand extracellular RNA transport and guide the design of therapeutic RNAs to maximize their incorporation in EVs towards targeted personalized medicine.

## Introduction

Extracellular vesicles (EVs) are membrane-bound vesicles that are secreted by most cell types (Colombo et al., 2014; Yáñez-Mó et al., 2015), ranging in size from small exosomes (∼30-100 nm) to larger microvesicles and apoptotic bodies (up to 1000 nm or more) (Raposo and Stoorvogel, 2013). These vesicles originate through distinct biogenesis pathways, carry diverse molecular cargo—including proteins, lipids, DNA, and various types of RNA—and perform a variety of biological functions (Chen et al., 2022; Payandeh et al., 2024). Once thought to be cellular “garbage bags,” EVs are now recognized as central mediators of intercellular communication in both physiological and pathological contexts, contributing to tissue homeostasis (Stratman et al., 2022; Tkach and Théry, 2016),(Stratman et al., 2022). Pathological conditions, such as cancer, can alter the composition of EV cargo, suggesting a highly regulated and selective mechanism of cargo sorting, though the underlying mechanisms remain incompletely understood (Abels and Breakefield, 2016; Gangoda et al., 2015; Miceli et al., 2024). Many of the effects mediated by EVs are attributed to their RNA cargo (Li et al., 2021). While microRNAs (miRNAs) have been extensively studied, other RNA species—including mRNAs, long non-coding RNAs (lncRNAs), and, more recently, circular RNAs (circRNAs)—are also selectively packaged into EVs (Wang et al., 2019). CircRNAs are covalently closed circular RNA molecules generated by back-splicing, representing approximately 1% of total cellular RNA, and are characterized by tissue-specific expression and high conservation across species (reviewed in (Hoque et al., 2023)). Their circular structure (i.e. the lack of free ends) confers resistance to exoribonucleases, rendering them considerably more stable than their linear counterparts highlighting their potential as excellent biomarker candidates (Li et al., 2015; Wang et al., 2020). Recent studies have reported the selective packaging of circRNAs into EVs, implicating them in intercellular communication and highlighting their role as promising non-invasive liquid biopsy biomarkers (Hon et al., 2019; Wang et al., 2019). However, the mechanistic details of circRNA packaging into EVs remain poorly understood, and comprehensive studies addressing this process are still lacking (Zhang et al., 2019).

Historically, RNA biology has focused on primary sequence (the order of nucleotides); however, it is now recognized that all RNAs adopt complex secondary and tertiary structures—including stem-loops, multi-helix junctions, and pseudoknots—that determine their three-dimensional shape and cellular fate (Assmann et al., 2023; Cao et al., 2024). RNA folding is a dynamic process which is influenced by intrinsic factors such as size, and nucleotide order and composition as well as extrinsic factors such as RNA-binding proteins (RBPs) and ionic concentration (Bushhouse et al., 2022). Intrinsic features including size, GC content and nucleotide order can be used to predict the thermodynamic free energy (ΔG) of folded structures which, along with other structure-related features, inform predictions of RNA structuredness and overall shape (Bellaousov et al., 2013; Busa et al., 2021; Lorenz et al., 2011; Low and Weeks, 2010; Radecki et al., 2018; Zuker, 2003). Advances in high-throughput RNA structure profiling and computational modeling have enabled transcriptome-wide characterization of RNA 3D structure (reviewed in (Strobel et al., 2018)),(Siegfried et al., 2014; Spitale et al., 2015; Tomezsko et al., 2021; Watters et al., 2018; Zhang et al., 2022; Zhu et al., 2025). While experimental methods are labor-intensive, they are essential for generating training data for computational tools. Despite the emerging nature of RNA structural biology, it is clear that RNA molecules exhibit substantial structural heterogeneity, and their structure significantly impacts function. For example, highly structured linear and circular RNAs are targeted by specific decay pathways, resulting in shorter half-lives (Fischer et al., 2020; Guo et al., 2020). Recent technologies have advanced our understanding of how RNA folding affects biology, particularly in mediating interactions with other RNAs and proteins (Chekulaeva, 2024; Dominguez et al., 2018). However, the role of RNA structure in intercellular transport, especially via EVs, remains largely unexplored, representing a critical knowledge gap for understanding RNA transport mechanisms.

Insights into the packaging of RNA cargo into EVs can be drawn from analogous systems, such as the selective packaging of viral genomes in positive-strand RNA viruses. Viruses efficiently select and package their own genomes amidst abundant host RNA, a process thought to be facilitated by the compact folding of viral RNAs, allowing them to fit within the limited space of the viral capsid (Ben-Shaul and Gelbart, 2015; Shrestha et al., 2018). Similarly, selective packaging of RNA cargo of varying sizes into EVs—which can be as small as 50 nm— may depend on structural features (Nolte-’t Hoen et al., 2016). Highly structured and compact RNAs may instead be sequestered in the cell or targeted for decay (Fischer et al., 2020). Notably, recent work from our group showed that RNAs selectively packaged into EVs exhibit distinct structuredness profiles compared to those retained in the cell (Abdelgawad et al., 2025). While advanced experimental techniques such as small angle X-ray scattering (SAXS) and atomic force microscopy (AFM) can assess RNA structure (Incarnato et al., 2018), secondary structure features—including minimum free energy (MFE), number of stems (SL), number of multi-loops of order three or higher (ML3), number of paired bases (BP), 90th percentile of base-pair distances (BP90), and maximum ladder distance (MLD)—can be used to predict RNA compactness and shape computationally (Danaee et al., 2018; Lorenz et al., 2011; Yoffe et al., 2008). For example, MLD measures the length of the longest uninterrupted path of non-crossing base pairs in the RNA’s secondary structure, analogous to the rungs of a ladder. Generally, RNAs with lower MFE and MLD values are more compact (Wayment-Steele et al., 2021). These features, calculated using prediction tools such as RNAfold, have been shown to be highly correlated with experimental measurements (Lorenz et al., 2011; Zuker, 2003) (Borodavka et al., 2016).

In this study, we built on our recent work (Abdelgawad et al., 2025) with a goal to predict RNA enrichment in EVs by analyzing cis elements—specifically, sequence and structural features—of both mRNAs and circRNAs. These features include size, GC content, nucleotide frequencies, structural characteristics, and coding probability for circRNAs. We also incorporated information on RNA-binding protein (RBP) interactions using both database-derived motifs and k-mer representations. After evaluating various models, we identified the best-performing approach for each RNA type, achieving an AUC of 0.9 and 0.77 for circRNAs and mRNAs, respectively. We subsequently developed a web-accessible tool, EVRCEPT, which enables researchers to estimate the likelihood of RNA enrichment in EVs. This tool may facilitate the identification of RNAs likely to be enriched in EVs, which could inform future studies on RNA therapeutic targets for human diseases.

## Material and Methods

### Dataset generation and classification

RNA-Seq dataset was downloaded from Gene Expression Omnibus (GSE279376) and was analyzed as previously mentioned except for using Gencode (v47) annotations for analyzing control samples (mRNAs) (Abdelgawad et al., 2025). Differential expression analysis was performed using DESeq2 (v1.42.1) R package with default parameters (Love et al., 2014) in R (v4.3.3). RNAs were labelled as follows: (1) EVs-enriched (EVE) if their fold change > 0 and their adjusted p value (FDR) < 0.05 or 0.2, for mRNAs or circRNAs, respectively. (2) cell-retained (CR) if their fold change < 0 and their adjusted p value (FDR) < 0.05 or 0.2, for mRNAs or circRNAs, respectively. (3) not significantly different (NS) for all remaining RNAs. Both mRNA and circRNA datasets were randomly split into 80% for training and 20% for testing.

### Statistical analysis

Pairwise correlation of features were computed using Spearman’s rank correlation coefficient on all numeric values using pandas (v2.2.1). To assess the association between features and the binary outcome variable, a one-way analysis of variance (ANOVA) F-test was used in scikit-learn (v1.6.1).

### Database motif representation

To represent RNA-binding protein motifs, 2 approaches were used. For the database approach, RBP motifs were downloaded from ATtRACT database (Giudice et al., 2016). Motifs from non-human species were filtered out and duplicates were removed which resulted in final list of 2,806 RBP motifs. Each motif was scanned across all sequences, and a binary encoding scheme was applied denoting the motif’s presence or absence. The resulting binary feature matrices was used as input to an autoencoder neural network. The autoencoder was constructed with two encoding layers (128 hidden units and 32 latent units with ReLU activation) and two decoding layers (128 hidden units with ReLU activation, followed by a sigmoid output layer matching the input dimension). The latent layer size of 32 was chosen to provide a compact representation while retaining relevant motif information. Training was performed for 50 epochs with a batch size of 32, using the Adam optimizer and binary cross-entropy loss. A 10% validation split from the training data was used for monitoring convergence. After training, the encoder module alone was applied to the binary motif matrices, producing 32-dimensional embeddings for both the training and test datasets. These embeddings served as reduced-dimension feature inputs for downstream models.

### K-mer motif representation

For the k-mer approach, k-mer features were generated using a bag-of-ngrams approach. K-mers of length 3 to 8 were selected to balance the ability to capture both short and moderately long sequence motifs while controlling feature dimensionality and computation cost. Feature extraction was implemented with CountVectorizer in scikit-learn (v1.6.1) producing sparse count matrices representing k-mer frequencies for each sequence. Sparse count matrices were then reduced in dimensionality using non-negative matrix factorization (NMF) with 32 components, initialized with the NNDSVD method and a maximum of 1000 iterations. The NMF transformation produced dense feature representations for both training and test sequences which were then used as inputs for downstream models.

#### Classification models

Features were standardized to zero mean and unit variance and were tested on 5 models: XGBoost, logistic regression, feedforward neural networks, random forests, and naïve Bayes. All models used the same training and test datasets, with the target variable encoded into binary labels. Both motif and k-mer representations were tested on all models. XGBoost was implemented with 300 trees, and maximum depth of 4 and learning rate of 0.05. Neural network was constructed with three hidden layers of 64, 32, and 16 neurons with ReLU activation, and dropout layers (rate 0.3) to reduce overfitting. The output layer consisted of a single neuron with sigmoid activation. The model was trained with binary cross-entropy loss and the Adam optimizer for 25 epochs. A random forest classifier was trained using 40 decision trees with a maximum depth of 3. For neural network and random forests models, class weights were used to account for class imbalance.

### Performance evaluation

Models’ performance was evaluated using several metrics including accuracy, balanced accuracy, precision, recall, F1 score and are under the receiver operating characteristic curve (AUC) as shown by the following equations:

1. 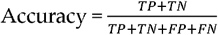
2. 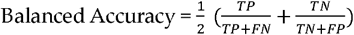
3. 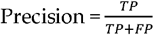
4. 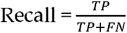
5. 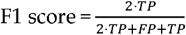
6. 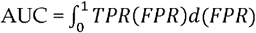 Where 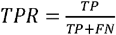 and 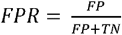where TP and TN denote correctly classified positive and negative samples, while FP and FN denote misclassified positive and negative samples, respectively. These measures account for possible imbalance between positive and negative classes.

### Software availability

Models were implemented in python 3 using Keras (v3.10.0), scikit-learn (v1.6.1) and xgboost (v3.0.4) packages. EVRCEPT was implemented using Shiny for Python (v1.4.0) and deployed on Shiny Server (v1.5.22.1017) and is available at https://euler.dbi.udel.edu/evrcept.

## Results

We utilized RNA-Seq dataset of mRNAs and circRNAs from DLD-1 cell and EV fractions to identify mRNAs and circRNAs (Abdelgawad et al., 2025). Identified RNAs were then classified into three categories: (1) significantly enriched in EVs (EVE), (2) significantly retained in the cell (CR), and (3) RNAs with no significant enrichment in either compartment (NS). To build a model predicting RNA enrichment into EVs, we used cis elements such as sequence and structural features, as well as information on potential RBP interactions. Most RNAs identified fell into the NS category (Figure 1A and Figure 2A). To focus on identifying RNAs likely to be enriched in EVs (EVE) and to simplify the prediction to a binary classification problem, we combined the CR and NS categories into a single group (NS)

**Figure 1:**
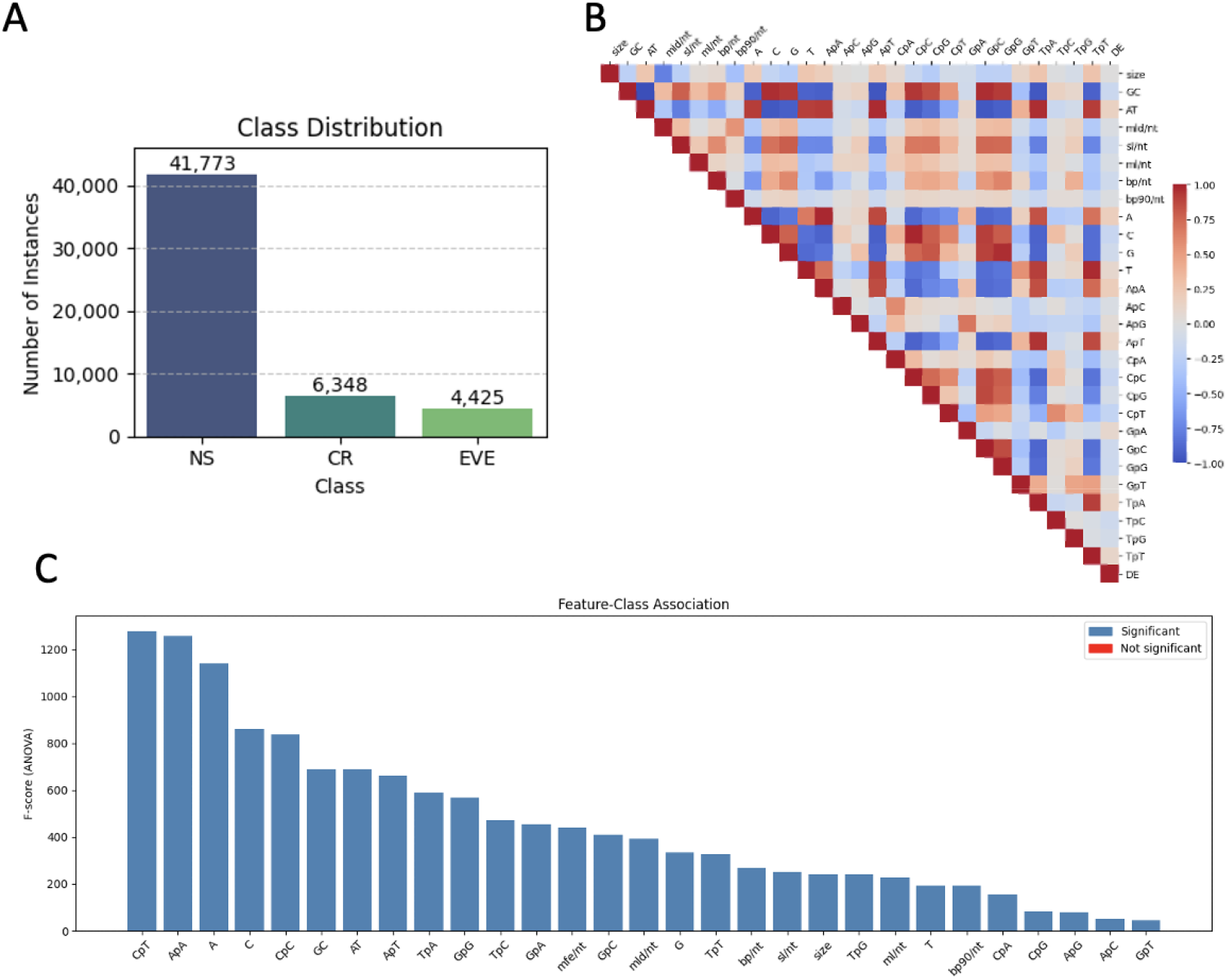
Feature analysis of mRNA data. (A) Label counts in mRNA dataset. (B) Spearman correlation matrix of pairwise correlation among numerical features as well as the target output used in the mRNA model. Only the upper triangle and diagonal are displayed. Color intensity indicates the strength and direction of correlation (red = positive, blue = negative) (C) ANOVA F-scores showing the strength of association between each feature and class labels in the mRNA model. Features colored in blue are statistically significant (p□<□0.05), while red indicates non-significant features.

**Figure 2:**
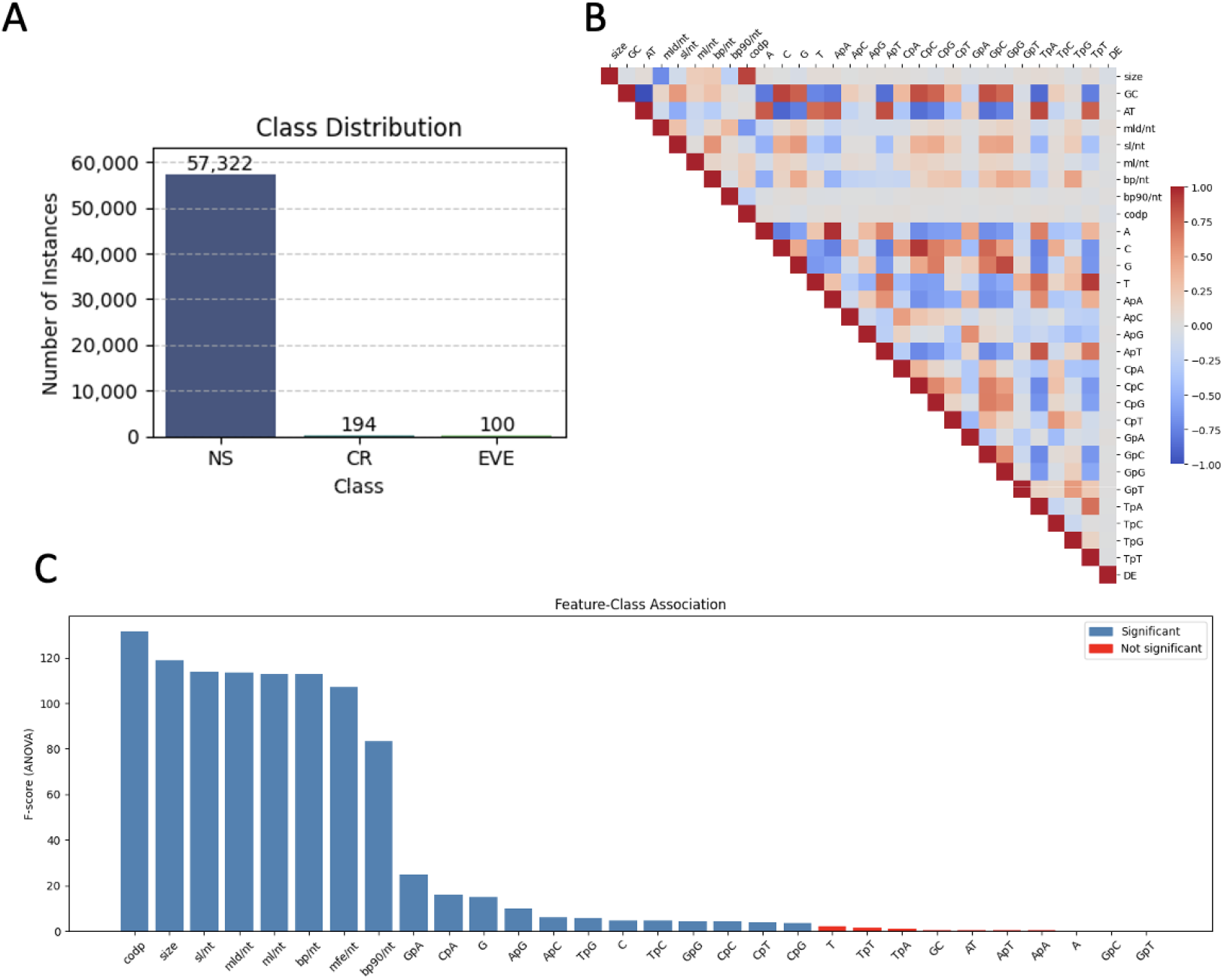
Feature analysis of circRNA data. (A) Label counts in circRNA dataset. (B) Spearman correlation matrix of pairwise correlation among numerical features as well as the target output used in the circRNA model. Only the upper triangle and diagonal are displayed. Color intensity indicates the strength and direction of correlation (red = positive, blue = negative) (C) ANOVA F-scores showing the strength of association between each feature and class labels in the circRNA model. Features colored in blue are statistically significant (p□<□0.05), while red indicates non-significant features.

We selected several sequence features for our model, including size, secondary structure parameters, nucleotide and dinucleotide frequencies, and coding probability for circRNAs (CODP). Six parameters were used to represent secondary structure: minimum free energy (MFE), maximum ladder distance (MLD), number of stems (SL), number of multi-loops of order three or higher (ML), number of base pairs (BP), and the 90th percentile of base pair distance (BP90). We examined the correlation of these features with RNA enrichment in EVs for both mRNA and circRNA datasets (Figure 1B and Figure 2B) and assessed feature importance using ANOVA F-scores (Figure 1C and Figure 2C). All features in the mRNA dataset and most in the circRNA dataset were found to be important.

Next, we incorporated information on cis factors representing RNA-binding protein (RBP) interactions, which involve recognition and binding to specific motifs within RNA. We considered two approaches: (1) utilizing RBP motifs from public datasets such as ATtRACT to calculate motif occurrences (Giudice et al., 2016). Although this database contains over 2,800 motifs, it represent only 179 RBPs which is less than a tenth of the estimated number of RBPs in human (Gerstberger et al., 2014; Qin et al., 2020). (2) we calculated the occurrence of all k-mer motifs (for k ranging from 3 to 8) in RNA sequences. We compared model performance using both of these approaches.

For the database approach, we encoded all motifs from the ATtRACT database (Giudice et al., 2016) as binary features, noting that many motifs had low occurrence in our dataset, particularly in the circRNA dataset (Figure 3A and Figure 4A). To reduce the high dimensionality (2,884 features), we trained an autoencoder on the binary motif embeddings, reducing them into a 32-dimensional representation. These compressed motif features were then combined with other sequence-based features (size, secondary structure, nucleotide, and dinucleotide frequencies) and were given as input to the model.

**Figure 3:**
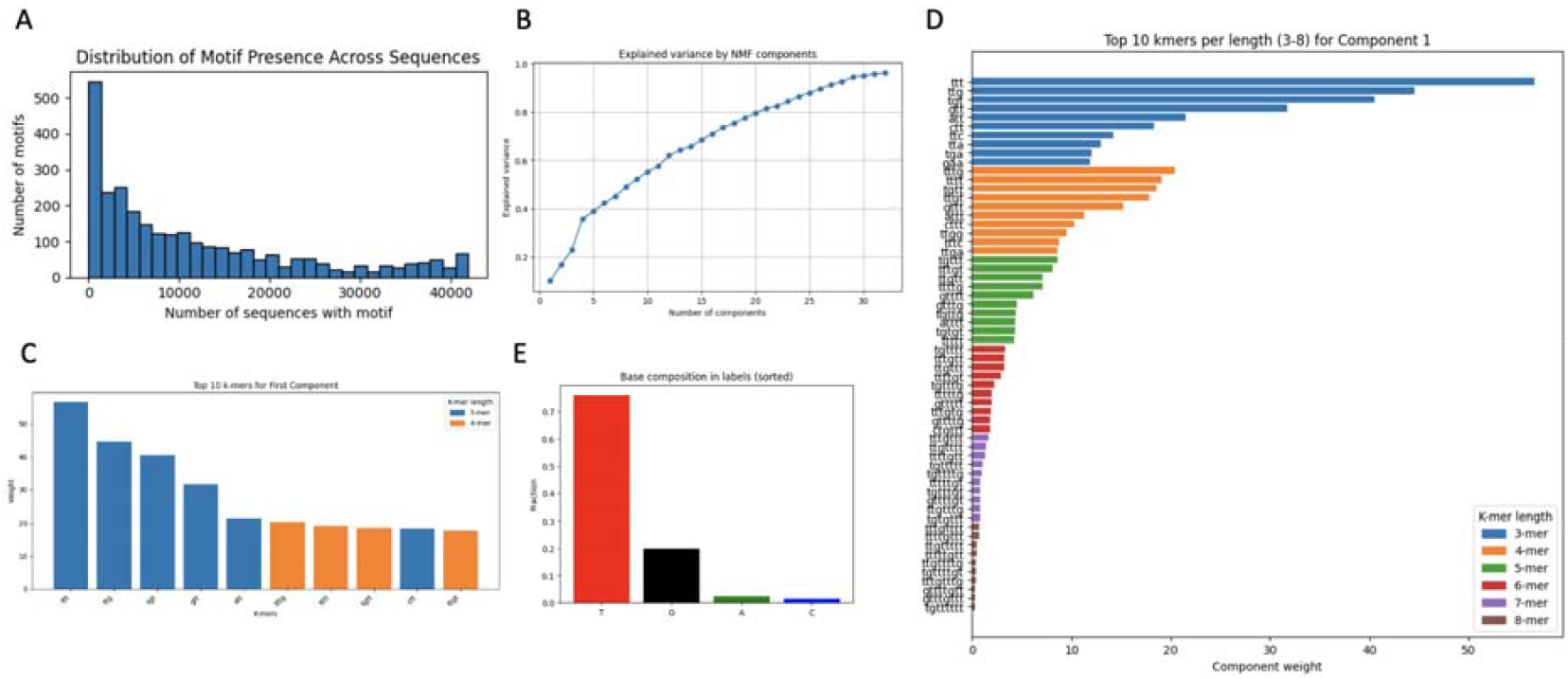
Sequence motif analysis of mRNA data. (A) Distribution of ATtRACT motifs in mRNA dataset. (B) Cumulative explained variance of the mRNA k-mer count matrix for the first 32 NMF components. (C) Top 10 k-mers by weight in the first NMF component for the mRNA dataset. (D) Top 10 k-mers by weight in the first NMF component for k3-k8 for the mRNA dataset. (E) Base composition of top 10 k3-k8 k-mers in the first NMF component.

**Figure 4:**
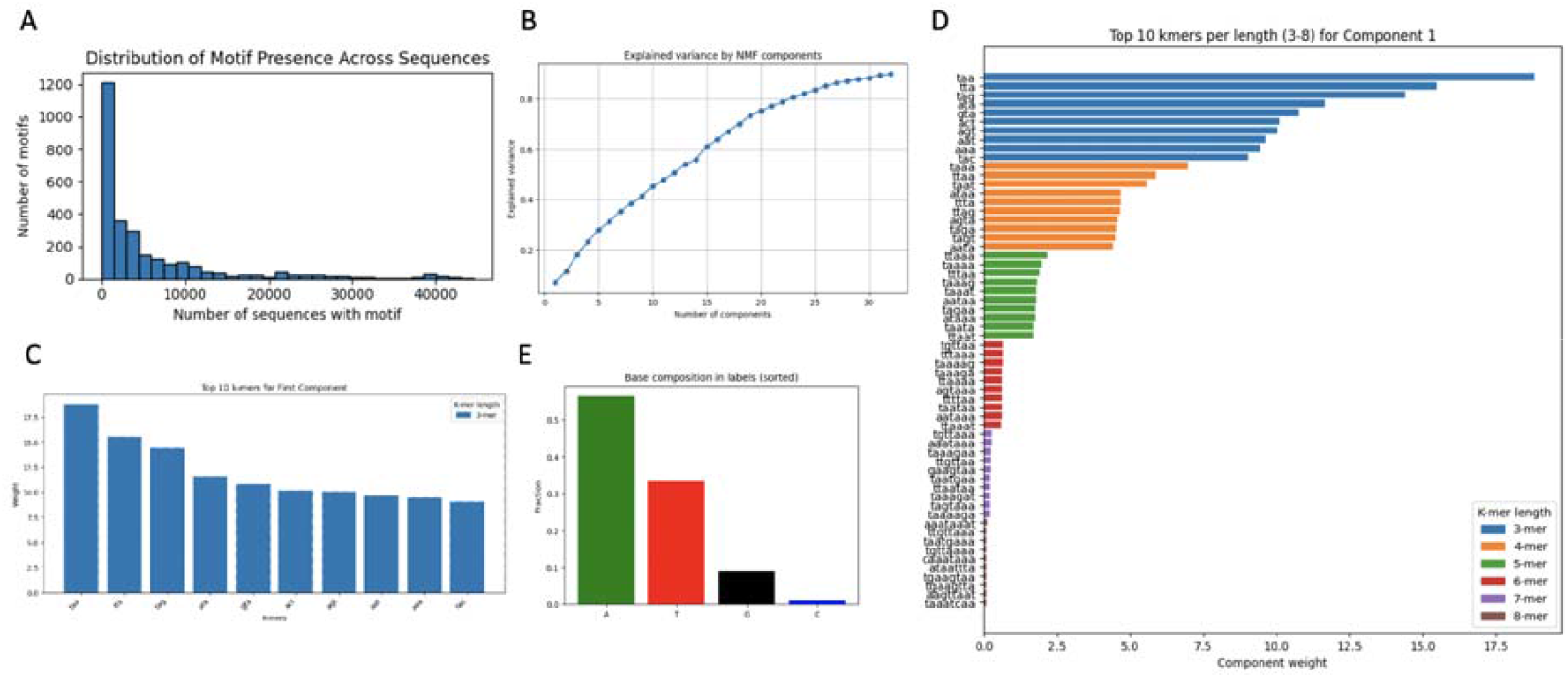
Sequence motif analysis of circRNA data. (A) Distribution of ATtRACT motifs in circRNA

For the k-mer approach, we calculated the occurrence of all 3-mers through 8-mers amounting to 87,360 features. To reduce dimensionality and identify latent sequence patterns, we applied non-negative matrix factorization (NMF) to the k-mer count matrices and selected the first 32 components for inclusion in the model alongside sequence and structure-based features. The cumulative explained variance for these components was approximately 0.90 for circRNAs and 0.96 for mRNAs (Figure 3B and Figure 4B). The top 10 motif contributions in the first NMF component were dominated by 3-mers since they occur more frequently than higher-order k-mers (Figure 3C and Figure 4C). Thus, we further analyzed the top 10 motif contributions for all k-mers (k3–k8) (Figure 3D and Figure 4D) and found that AT-rich motifs were predominant for circRNAs, while TG-rich motifs were most prominent for mRNAs (Figure 3E and Figure 4E).

We tested 5 different models including extreme gradient boosting (XGBoost) logistic regression (LR), and deep neural network (NN), random forests (RF), and naïve bayes (NB). We assessed the performance of these models using 6 metrics including overall accuracy, balanced accuracy, precision, recall, F1 score and AUC (Table 1).

**Table 1.** Performance of 5 models tested on circRNA (top) and mRNA (bottom) datasets.

| Metric | XGB |  | LR |  | NN |  | RF |  | NB |  |
| --- | --- | --- | --- | --- | --- | --- | --- | --- | --- | --- |
|  | Motifs | Kmers | Motifs | Kmers | Motifs | Kmers | Motifs | Kmers | Motifs | Kmers |
| Accuracy | **0.998** | 0.998 | 0.784 | 0.768 | 0.918 | 0.735 | 0.770 | 0.756 | 0.436 | 0.537 |
| Balanced Accuracy | 0.500 | 0.500 | 0.740 | 0.754 | 0.590 | 0.629 | **0.885** | 0.856 | 0.718 | 0.768 |
| Precision | 0.996 | 0.996 | 0.997 | 0.997 | 0.996 | 0.997 | **0.998** | 0.998 | 0.998 | 0.998 |
| Recall | **0.998** | 0.998 | 0.784 | 0.768 | 0.918 | 0.735 | 0.770 | 0.756 | 0.436 | 0.537 |
| F1 score | **0.997** | 0.997 | 0.877 | 0.867 | 0.956 | 0.845 | 0.868 | 0.859 | 0.605 | 0.697 |
| AUC | 0.885 | 0.893 | 0.835 | 0.835 | 0.790 | 0.763 | **0.904** | 0.896 | 0.868 | 0.867 |
|  | Motifs | Kmers | Motifs | Kmers | Motifs | Kmers | Motifs | Kmers | Motifs | Kmers |
| Accuracy | **0.913** | 0.913 | 0.668 | 0.680 | 0.656 | 0.675 | 0.667 | 0.657 | 0.136 | 0.708 |
| Balanced Accuracy | 0.503 | 0.503 | 0.701 | **0.705** | 0.693 | 0.704 | 0.677 | 0.679 | 0.517 | 0.663 |
| Precision | 0.881 | **0.906** | 0.895 | 0.895 | 0.893 | 0.895 | 0.888 | 0.889 | 0.888 | 0.882 |
| Recall | **0.913** | 0.913 | 0.668 | 0.680 | 0.656 | 0.675 | 0.667 | 0.657 | 0.136 | 0.708 |
| F1 score | **0.872** | 0.872 | 0.740 | 0.749 | 0.731 | 0.745 | 0.739 | 0.731 | 0.110 | 0.769 |
| AUC | 0.759 | 0.766 | 0.757 | 0.764 | 0.757 | **0.770** | 0.740 | 0.741 | 0.702 | 0.712 |

For circRNAs, the RBP motif representation performed slightly better than the k-mer representation, with the random forest model achieving the highest AUC score of 0.904 (Figure 5A). Models trained on the mRNA dataset had generally lower performance as compared to those for circRNAs. We found that RBP motif representations performed better in some metrics while k-mer representation performed better in others. Overall, neural network model with k-mer representation achieved the highest AUC score of 0.77 (Figure 5B).

**Figure 5:**
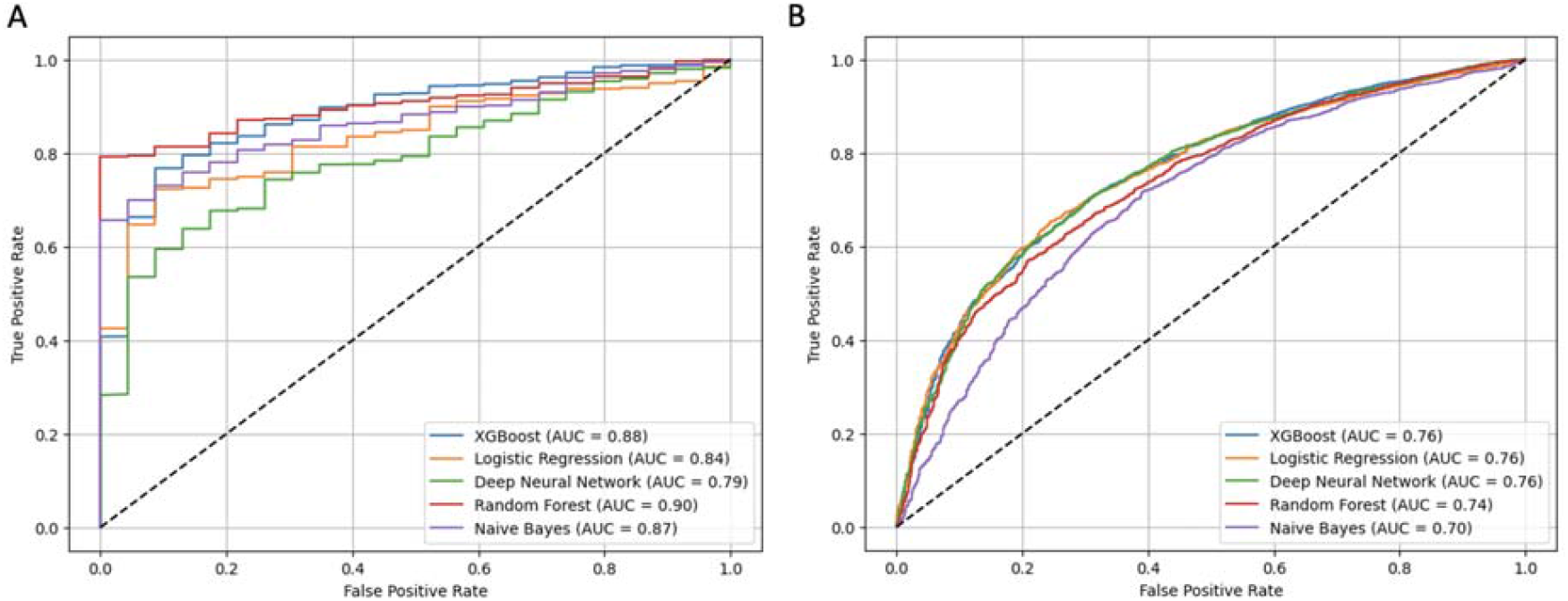
Performance evaluation of RNA enrichment prediction models. (A, B) ROC curves of 5 models with RBP motif representation tested on circRNA (A) and mRNA (B) datasets, respectively. AUC for each model is shown in brackets.

We, then, used this information to develop a web accessible tool (called EVRCEPT) to help guide researchers in predicting RNAs enrichment into EVs. The tool has a simple, user-friendly interface and only requires an RNA sequence as input (Figure 6A). The user can paste a sequence in either DNA or RNA format and select the type of RNA (mRNA or circRNA) to predict its enrichment in EVs. The model calculates an enrichment score which ranges from 0 (highly likely to be retained in the cell) to 1 (highly likely to be enriched in EVs) (Figure 6B-C). The input sequence is restricted to a size range of 10 nt to 32,767 nt. Using this tool, researchers can easily compare different RNA variants to see which is more likely to be enriched in EVs. We utilized the random forest model with RBP motif representation for circRNAs and neural network model with k-mer representation for mRNAs. These models were only trained on human transcriptome data but as we include more datasets from other species, the user will be able select a species for a given sequence. Once the user initiates the prediction, the tool will extract all relevant sequence and structural features from the input sequence, runs the predictions and then provides a breakdown of those features under the “Features” tab (Figure 7A) The tool will also quantify the motifs of all RBPs in the ATtRACT database and show the number of distinct motifs for each RBP as well as the total number of motifs present in the input RNA in both a tabular and graphical representations which be shown under the “Motifs” tab (Figure 7B). The computation time depends on the size of the input but for a sequence size of 1 kb, the tool returns results within a few seconds.

**Figure 6:**
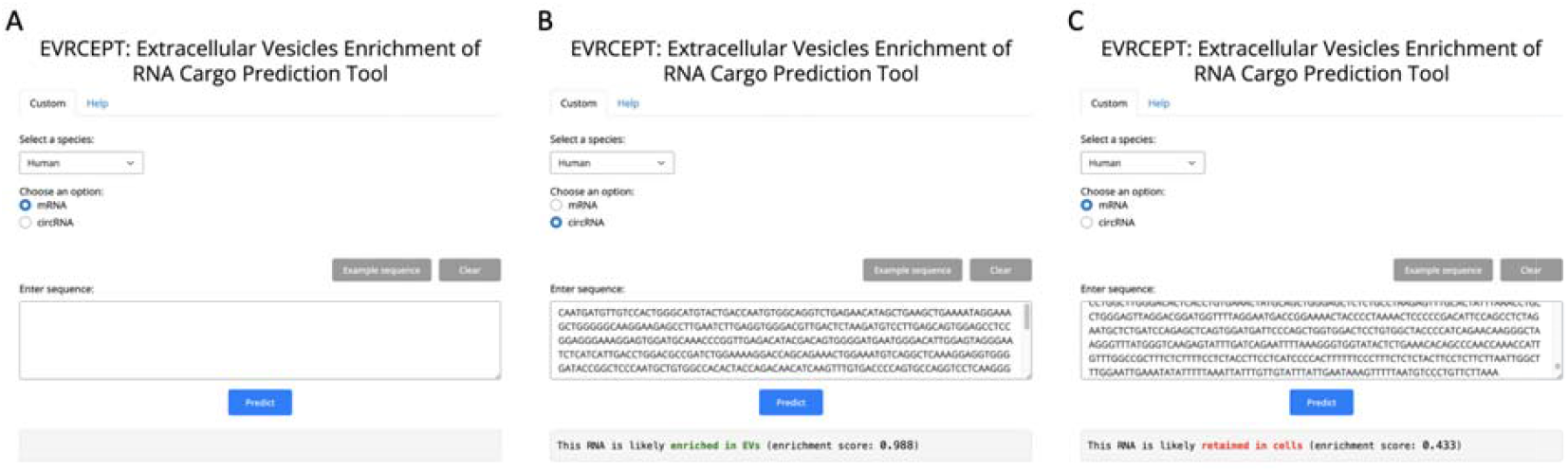
EVs RNA Cargo Enrichment Prediction Tool (EVRCEPT) user interface. (A) EVRCEPT simple and user-friendly interface. The user can input a sequence in DNA or RNA format and select the RNA type (mRNA or circRNA). (B, C) Predicted enrichment of 2 example RNA sequences that are likely to be enriched in EVs (B) or retained in the cell (C) with enrichment score shown in parentheses.

**Figure 7:**
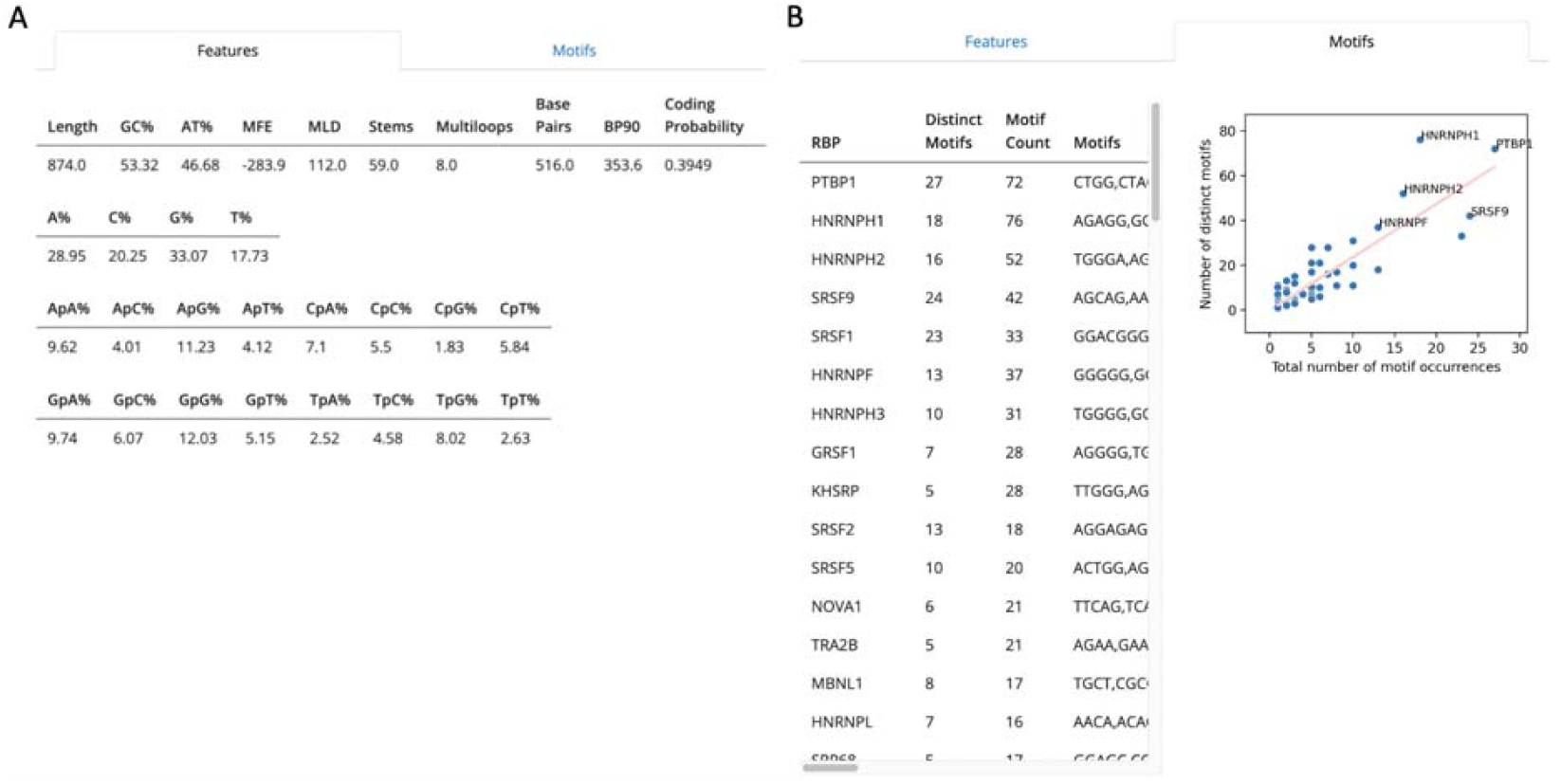
EVRCEPT output features. (A) Sequence and structural features calculated from the user sequence that were used as input to the model. (B) Tabular and graphical representations of the number of distinct and total motifs for each RBP from the ATtRACT database that are present in the input sequence.

## Discussion

In this study, we developed a robust computational tool to predict RNA enrichment in EVs which builds on previous studies that explored the mechanisms governing selective RNA packaging. We previously identified the sequence and structural features of RNAs that are correlated with higher enrichment in EVs. Specifically, we showed that RNAs that are enriched in EVs tend to be less structured and less compact than RNAs retained in the cell (Abdelgawad et al., 2025). Interestingly, features derived from secondary structure, such as MLD and MFE, were strong predictors of RNA enrichment especially for circRNAs. This might be because a high MLD indicating a more linear or extended RNA structures may be favored for packaging via facilitating recognition by the sorting machinery. Furthermore, NMF analysis highlighted distinct k-mer patterns which had higher proportions of AT content. Despite the similitude between RNA cargo packaging in EVs and viral RNA packaging, there seem to be some differences. Viral genomes are on the order of several thousand nucleotides where specific structural features are essential for efficient fitting into a constrained space. On the other hand, most RNA cargo in small EVs are typically a few hundred nucleotides long. However, the exact structural requirements warrant further investigation. We have also incorporated RBPs’ interaction into the model using two different approaches. Overall, database and k-mer approaches showed slight differences in performance across all models tested. We, ultimately, selected the one that achieved the higher performance to be deployed on our webtool. Databases such as EVLncRNAs incorporates experimentally validated lncRNAs in EVs which underscores the growing recognition of these molecules as critical EV cargo and further highlights the emerging need for tools like EVRCEPT to accurately predict their packaging (Zhou et al., 2024). The predictive capability of our tool may have substantial translational potential; by identifying EVs-enriched RNAs, EVRCEPT could support the discovery of candidate non-invasive liquid biopsy biomarkers for early disease detection and monitoring. Moreover, this tool may assist in the design of therapeutic RNAs that are engineered for optimal EV packaging.

Although this study provides a robust tool for EV RNA cargo prediction, several limitations should be acknowledged. First, the model relies primarily on predicted RNA secondary structure, which is constrained by the accuracy and size limitations of current prediction tools. For example, EVRCEPT has a size limit of 32,767 nt due to RNAfold input constraints (Lorenz et al., 2011). Ultimately, understanding the mechanistic details of RNA packaging will require accurate 3D structure predictions and high-resolution experimental data to train and validate such tools. Despite this limitation, however, our study offers the first comprehensive computational link between specific RNA structural features, sequence motifs, and selective EV packaging, providing a foundation for future mechanistic studies. Second, the predictive models are limited by the size and diversity of the training data. Although we identified tens of thousands of mRNAs and circRNAs, the training data are derived from a single RNA-Seq dataset, which may introduce cell type bias. Future research should incorporate more datasets, particularly for circRNAs, which often require specialized processing such as RNase R treatment. Expanding the dataset will increase the number of positive-class RNAs, alleviate class imbalance, and improve the overall classification performance.

We plan to maintain and update EVRCEPT in the foreseeable future in two main areas: model expansion and refinement and enhanced functionality of the webtool. First, we will expand the scope of EVRCEPT by including new predictive models for other types of RNAs, such as lncRNAs, which are emerging as significant EV cargo. We will continually improve the existing models by adding new experimental data as it becomes available and incorporating novel sequence and structural features identified through ongoing bioinformatics research. We will maintain the model’s relevance by consistently testing and updating predictions on new public EV RNA-seq databases as they are released or updated. Second, we will add new functionalities to streamline data input. This includes linking EVRCEPT with specialized circRNA databases, such as CircInteractome (Dudekula et al., 2016), allowing the user to simply submit a circRNA ID to automatically retrieve the sequence required for prediction. Similarly, we plan to enable the user to submit an ENSEMBL transcript ID for mRNAs. Finally, to broaden the tool’s utility in preclinical research, we plan to develop dedicated predictive models for EV cargo enrichment in other species such as mouse (*Mus musculus*). In summary, EVRCEPT is a valuable tool that may support efforts in biomarker discovery and engineering of therapeutic RNAs for EV-based delivery.

## Data Availability Statement

The source code is available at http://github.com/ijyrem/evrcept-model.

## Acknowledgments

We would like to thank the members of Batish Laboratory for their discussion and support. We also acknowledge Omar Ahmed for his help in building and validating the models. We thank the support from the University of Delaware Bioinformatics Data Science Core Facility (RRID:SCR_017696) including use of the BIOMIX and BioStore computational resources that was made possible through funding from Delaware INBRE (NIGMS P20GM103446), NIH Shared Instrumentation Grant (NIH S10OD028725), the State of Delaware, and the Delaware Biotechnology Institute.

## Ethics approval statement

N/A

## Patient consent statement

N/A

## Permission to reproduce material from other sources

N/A

## Clinical trial registration

N/A

## Author Contributions

AA: Investigation, data curation, analysis, methodology, article writing, review, and editing. JB: Methodology, review, and editing. SP: Methodology, review, and editing. VP: Funding acquisition, review, editing. MB: Conceptualization, supervision, analysis, funding acquisition, project administration, article writing, review, and editing.

